# Partner fidelity and coevolution: Useful but not required for rapidly increased mutualistic benefits

**DOI:** 10.1101/2025.09.07.674547

**Authors:** Kayla S. Stoy, Dung Lac, William C. Ratcliff

## Abstract

Evolutionary theory predicts that specialization between mutualistic partners is beneficial, limiting conflict and increasing the opportunity to evolve cooperative benefits through sustained reciprocal selection. However, specialized mutualisms are relatively rare in nature. Few empirical studies have directly examined how multi-partner interactions affect mutualistic evolution, largely because tracking partner dynamics over evolutionary timescales is challenging in natural systems. We circumvent this constraint via experimental evolution with an engineered resource sharing mutualism in Baker’s yeast (*Saccharomyces cerevisiae*), which allows precise control of partner fidelity across generations. We compared high partner fidelity (consistent pairings) with low partner fidelity (temporally rotating partners) across 54 rounds of selection. High partner fidelity produced the strongest mutualistic benefits (7.2% increased growth over controls), while low partner fidelity yielded only modest benefits (2.2% increase). Time-shift experiments confirmed that while coevolution enhanced benefit evolution, it was not strictly necessary: benefits also evolved through one-sided adaptation. Genomic analyses revealed parallel evolution in amino acid metabolism and starvation response genes, with the slower-growing genotype driving most evolutionary change. Surprisingly, high-fidelity lineages evolved as generalists rather than specialists, cooperating equally well with all partners tested. Overall, these results demonstrate that partner fidelity facilitates the rapid evolution of mutualistic benefits but is not strictly required for mutualistic evolution.

## Introduction

Most organisms rely on mutualism – cooperative interactions between species where both receive a benefit (Bronstein 2015). Mutualism underpins organismal fitness and function (Baumann et al. 1995; Mueller and Gerardo 2002; Gerardo et al. 2020; Kaltenpoth and Flórez 2020a), contributes to species diversification (Weber and Agrawal 2014; Chomicki et al. 2019; Cornwallis et al. 2023; García-Lozano et al. 2024), and increases ecosystem biodiversity and stability (Bascompte et al. 2003; Hale et al. 2020). While nearly all organisms depend on another species for essential resources or services, participating in mutualism is expected to come at a cost (Sachs et al. 2004; Foster and Kokko 2006; Douglas 2008; Weyl et al. 2010; Frederickson et al. 2012; Masson et al. 2015). The energetic investment required to produce resources or provide services for a heterospecific partner may directly reduce fitness (Strauss et al. 2002; Frederickson et al. 2012). Consequently, mutualists should experience selection to act selfishly and cheat their partner by receiving benefits without reciprocating (Bronstein 2001).

Specialization can evolve as partners impose selection on one another to both limit cheating and increase benefit exchange (Parker 1995; Frank 1996; Douglas 1998; Moran and Wernegreen 2000; Thrall et al. 2007). Specifically, partners may limit cheating by preferentially interacting with and rewarding only beneficial partners and avoiding, or sanctioning, less cooperative partners, which can drive specialization between the most beneficial partner genotypes (Axelrod and Hamilton 1981; Noe and Hammerstein 1994; Frank 1996; Kiers et al. 2003; Sachs et al. 2004; Thrall et al. 2007; Leeks et al. 2019). Partner fidelity enhances opportunities for specialization by enabling partners to impose persistent selection on one another for cooperative traits and increased mutualistic benefit (Parker 1995; Thompson 2005). Mutualistic benefits may evolve most rapidly when both partners impose reciprocal selection on one another through the processes of coevolution, *i.e.*, reciprocal evolutionary change between species driven by natural selection (Thompson 1994, 2005). Consequently, coevolutionary specialization between mutualistic partners is thought to facilitate the evolution and persistence of mutualism, limiting conflict and increasing the opportunity to evolve cooperative benefits (Thompson 2005).

While notable examples of highly specialized interactions exist, mutualists are most often generalists that interact with multiple genetically diverse partners across space or time (Fisher et al. 2017; Batstone et al. 2018; Chomicki et al. 2020). Often generalist interactions are ancient (Hilário et al. 2011; Kikuchi et al. 2011; De La Peña et al. 2018) and obligate (Hilário et al. 2011; Hosokawa et al. 2016; Chomicki et al. 2020; Kaltenpoth and Flórez 2020b), indicating they are not simply transition states on the path to increased specialization. The long-term evolutionary consequences of the low partner fidelity associated with generalism remain poorly understood. It is unclear whether processes like coevolution that can drive benefit evolution in specialized mutualisms also underpin the stability and evolution of mutualistic benefits in interactions involving multiple partners (Hall et al. 2020). The low partner fidelity associated with generalism should limit sustained selection between partners, potentially inhibiting coevolution and impeding the evolution of increased mutualistic benefits. Yet the prevalence and apparent stability of generalist mutualisms suggest partners readily impose reciprocal selection on one another through which increased mutualistic benefits are likely to evolve. Evaluating how partner fidelity affects these fundamental evolutionary processes is important for understanding how and when mutualistic benefits can evolve and persist across the specialization-generalism spectrum.

We used experimental evolution to test how temporal variation in partner identity alters the evolution of mutualistic benefits. Specifically, we tested the hypothesis that low temporal fidelity between mutualistic partners limits opportunities for coevolution and constrains the evolution of mutualistic benefits. While generalism is common in extant mutualisms (Batstone et al. 2018; Chomicki et al. 2020; Lajoie and Parfrey 2022), isolating the effects of multi-partner interactions on mutualistic outcomes can be challenging in systems where the temporal sequence of interactions cannot be continuously monitored, and multiple selective processes operate simultaneously. Experimental evolution leveraging mutualistic model systems circumvents these limitations (Shapiro and Turner 2014; Turner et al. 2016; Hoang et al. 2016, 2021, 2022, 2024; Morran et al. 2016; Rafaluk-Mohr et al. 2018; Batstone et al. 2020; Vidal et al. 2020, 2024, 2025; Vidal and Segraves 2021; Debray et al. 2022; Ordovás-Montañés et al. 2022; Pauli et al. 2022; Stoy et al. 2023b). Using a genetically engineered model system of *Saccharomyces cerevisiae* strains that are obligately dependent on one another for the mutualistic cross-feeding of the amino acids tryptophan and leucine, we directly examined how temporal variation in partner identify alters mutualistic evolution.

In this study, we consider generalist mutualists as those interacting with multiple temporally variable partner strains. Across experimental evolution treatments, we manipulated the temporal fidelity of partner strains across experimental treatments (Figure 1). In the high partner fidelity treatment (HPF), we maintained consistent associations between specific pairs of partner strains over 54 rounds of selection. Co-passaging these strains created opportunities for pairwise coevolution. In the low partner fidelity treatments (LPF), we subjected focal mutualists to selection from partner strains that rotated through time. Here, partners still had opportunities for repeated interactions across experimental passages that could possibly permit diffuse coevolution (*i.e.*, coevolution between interacting guilds of species (Janzen 1980)). However, in the LPF treatments, the temporal rotation of partner strains meant focal mutualists were not consistently paired with the same partners, reducing the efficacy of selection between specific partner pairs and weakening coevolution compared to the HPF treatment. Within the control treatments, both partners evolved independently without selection for mutualism. We then tested the consequences of manipulating temporal partner fidelity by measuring mutualism fitness across experimental treatments. Overall, our results suggest that coevolution between partners evolved with high fidelity most rapidly enhances mutualistic benefits, but increased benefits can still evolve even when partner fidelity is low and coevolution does not occur.

**Figure 1:**
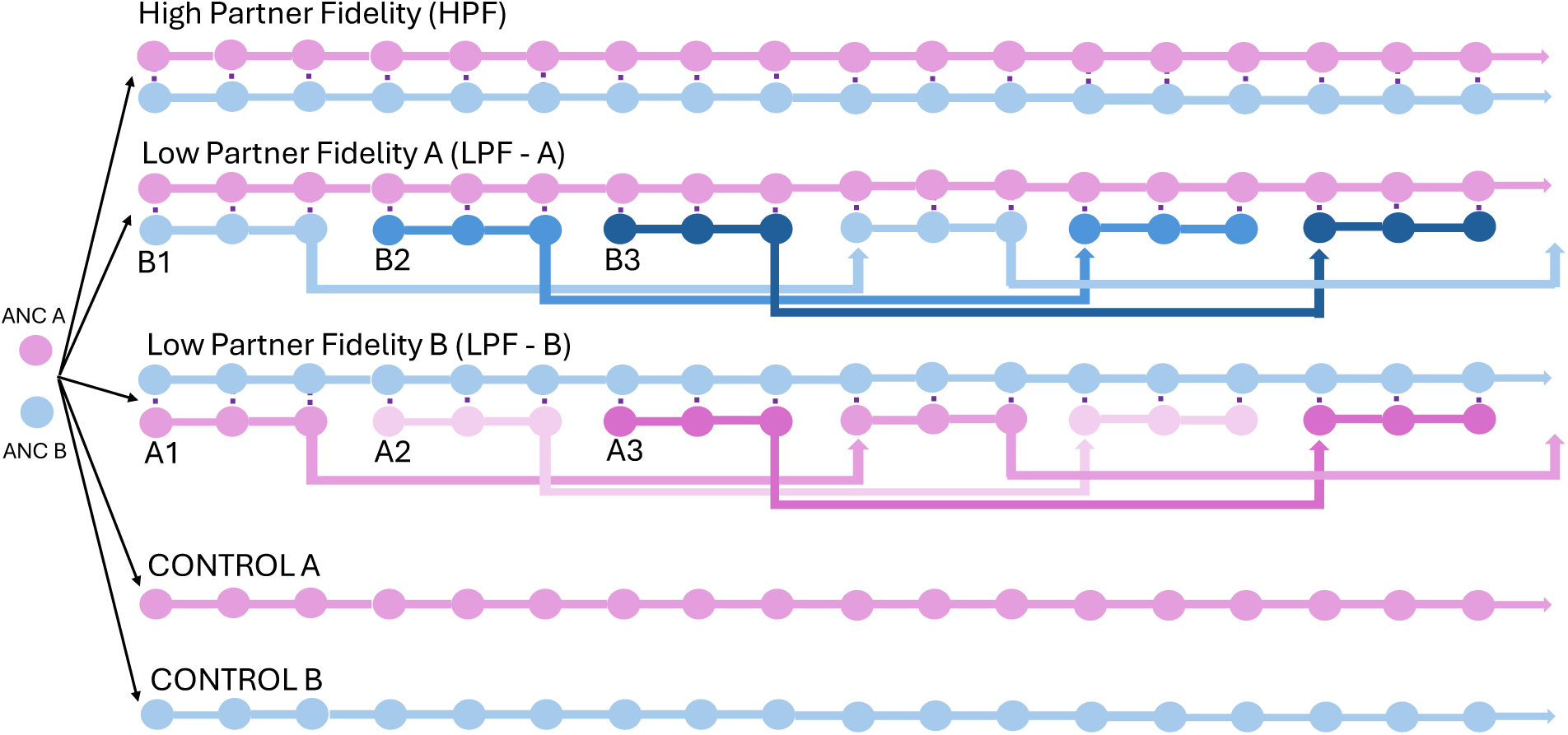
Experimental evolution design manipulating partner fidelity in mutualistic yeast. Partner A strains (Leu+Trp-; pink) and Partner B strains (Leu-Trp+; blue) began as isogenic clones and were subjected to five treatment conditions. In the <u>H</u>igh <u>P</u>artner <u>F</u>idelity treatment (HPF), Partner A and Partner B strains were strictly co-passaged throughout the experiment, permitting tight coevolution. In the <u>L</u>ow <u>P</u>artner <u>F</u>idelity A treatment (LPF-A), focal Partner A strains experienced selection from three different rotating Partner B strains (B1, B2, B3). In the <u>L</u>ow <u>P</u>artner <u>F</u>idelity B treatment (LPF-B), focal Partner B strains experienced selection from three different rotating Partner A strains (A1, A2, A3). Each rotating strain began as genetically identical clones and differences across strains resulted from the independent accumulation of *de novo* mutations. In both LPF treatments, rotating partners were co-passaged with the focal mutualist for three consecutive passages, stored at -80 °C, then reintroduced every tenth round of selection. This limited the efficacy of selection between specific partner pairs, but repeated interactions between partners across time permitted weak coevolution. Selection for mutualism in HPF and LPF treatments (five replicate populations each) occurred on SC-Leu-Trp plates, while control treatments (three replicate populations each) allowed Partner A and B to evolve independently on SC plates supplemented with leucine and tryptophan. All strains were haploid and obligately asexual. The experiment continued for 54 rounds of selection for focal mutualist evolution (18 rounds of selection per rotating partner). After 54 rounds of selection, we selected a single isolate of each strain from each replicate population, which served as the representative genotype for that replicate population for all growth assays and genomics.

## Methods

### Mutualistically cross-feeding yeast system

To generate a model system of cross-feeding mutualism, we modified strains of Baker’s yeast (*Saccharomyces cerevisiae*) described in Müller *et al*., 2014. Briefly, haploid, asexually reproducing Leu^+^Trp^-^and Leu^-^Trp^+^ strains were isogenically derived from *S. cerevisiae* strain W303. Strains have the following shared genetic background: MATa can1-100 hmlαΔ::BLE leu9Δ::KANMX6 prACT1yCerulean-tADH1@URA3. Amino acid overproduction results from feedback resistance mutations in *LEU4* and *TRP2* that make them insensitive to feedback in inhibition by leucine and tryptophan, respectively (Graf et al. 1993; Cavalieri et al. 1999). The Leu^+^Trp^-^ strain has the following unique modifications that make it express the yellow fluorescent protein mCitrine, overproduce leucine, auxotrophic for tryptophan, and resistant to natamycin (NAT): his3Δ::prACT1−ymCitrine− tADH1:HIS3MX6 LEU4^FBR^ trp2Δ::NATMX4. The Leu^-^Trp^+^ strain has the following unique modifications make it express the red fluorescent protein mCherry, auxotrophic for leucine, overproduce tryptophan, and resistant to hygromycin: his3Δ::prACT1−ymCherry−tADH1:HIS3MX6 leu4Δ::HPHMX4 TRP2^FBR^. Both strains fluorescently express the blue fluorescent protein yCerulean.

### Overview of Experimental Evolution

We hereafter refer to the Leu^+^Trp^-^ genotype as Partner A and the Leu^-^Trp^+^ genotype as Partner B for ease of communicating the experimental evolution design. We selected for mutualistic cross-feeding across three separate treatments (Figure 1). In the high partner fidelity (HPF) treatment, lineages of Partner A and Partner B were consistently co-passaged, resulting in high temporal fidelity. In the low partner fidelity (LPF) treatments, Partner A and Partner B were co-passaged with rotating partners, resulting in low temporal fidelity. We created two symmetric treatments, holding one “focal partner” fixed in each. In the low partner fidelity A (LPF-A) treatment, Partner A constantly evolved in response to three isogenic Rotating Partner B strains (B1, B2, and B3). In the low partner fidelity B (LPF-B) treatment, Partner B constantly evolved in response to three isogenic Rotating Partner A strains (A1, A2, and A3). Rotating Partner B and Rotating Partner A strains underwent three consecutive co-passages during each exposure to the constantly evolving LPF-A and LPF-B focal partners, and rotating partners were reintroduced every tenth round of selection (Figure 1). HPF, LPF-A, and LPF-B treatments were independently replicated five times. In Control A and Control B, lineages of Partner A and Partner B were passaged without exposure to one another on agar supplemented with all required amino acids. Control treatments were each independently replicated three times. A total of 54 experimental passages were conducted for each treatment. For the LPF treatments, this resulted in 18 total passages for each rotating partner.

### Transfer and selection for mutualism

To initiate experimental evolution each ancestor was streaked on synthetic complete (SC) agar plates [0.67% yeast nitrogenous base (w/v), 2% glucose (w/v), 0.087% amino acid mixture (w/v), 2% agar (w/v)] and grown at 30 °C for 48 hours. We selected a single colony forming unit (CFU) from each ancestor and separately inoculated them into 10mL SC liquid. Cultures were grown overnight at 30 °C with shaking at 250 RPM. Overnight cultures were used to initiate the experimental evolution of all replicate populations except those belonging to Rotating Partner A2, Rotating Partner A3, Rotating Partner B2, and Rotating Partner B3 (Figure 1). These replicate populations were also initiated as described above and used the same clonal ancestral stocks. However, replicate populations of Rotating A2/Rotating B2 were initiated prior to the fourth round of selection and Rotating A3/Rotating B3 populations were initiated prior to the seventh round of selection (Figure 1).

We removed SC media from cultured cells by washing 2mL of each liquid culture three times in 1mL 0.67% yeast nitrogenous base (YNB) and resuspended cells in YNB at a final volume of 500uL. We prepared aliquots for each replicate population by removing 150µL of washed culture and measuring OD600. To ensure equal starting conditions, aliquots were diluted to an OD600 of 1.0 in a final volume of 400 µL, and co-cultures were prepared by mixing Partner A and Partner B in equal volumetric ratios. We prepared 24-well selection plates by pipetting 2mL of SC agar lacking leucine and tryptophan (SC-Leu-Trp) into 17 wells (for mutualism selection) and 2mL of SC agar containing leucine and tryptophan into 6 wells (for controls). Co-cultures were seeded onto SC-Leu-Trp agar by placing 20µL drops in the center of each well. Controls were seeded by separately placing 20uL drops of washed and diluted cultures of each genotype onto the center of SC agar wells. Negative controls were used to test for cross-contamination and seeded by separately placing 20µL drops of each genotype onto SC-Leu-Trp wells. Plates were incubated at 30 °C. Ancestral genotypes required approximately four days for noticeable colony growth (Figure S1), so colonies were passaged to new agar plates every fifth day.

Cultures were consecutively passaged for three rounds of selection prior to introducing a new rotating partner (Figure 1). To do this, colonies were removed from agar using a sterile inoculating loop and suspended in 500uL YNB (6.7%). Cell suspensions were then vortexed, and 20µL of each suspension were seeded into new wells. In the Diffuse A and Diffuse B treatments, new rotating partners were introduced every three passages (Figure 1), which required strain separation using antibiotics. To minimize variation, we exposed all treatments to antibiotic selection. At the end of every third round of selection, colonies were removed from agar using an inoculating loop, suspended into 1mL YNB, and mixed by vortexing. To isolate Partner A, 50µL of each suspension was seeded into 10mL liquid SC+Hygromycin (800µg/mL), and to isolate Partner B 50µL of each cell suspension was seeded into 10mL liquid SC+Natamycin (400µg/mL). Cultures grew in antibiotic selection media at 30 °C with shaking for 24 hours.

Fifty microliters of each culture were exposed to a second round of antibiotic selection to dilute any spontaneous antibiotic-resistant mutants by transferring 50µL of each culture to a fresh 10mL of antibiotic selection media. We verified successful strain separation and screened for antibiotic resistant mutants by washing 2mL aliquots of each culture three times in YNB and separately plating 20µL of each washed culture onto SC-Leu-Trp and confirming a lack of growth. We performed this screening protocol following each round of antibiotic selection (every three passages).

Glycerol stocks of each replicate population were prepared from washed cultures every three rounds of selection and stored at -80 °C. To ensure reintroductions of Rotating Partner A and Rotating Partner B captured the population variation that arose during previous exposures, we prepared two single use 250µL aliquots (1 for reintroduction + 1 backup) of each washed rotating partner population. To reintroduce each rotating partner, one vial for each population was thawed, and populations were revived by suspending the entire sample into 10mL SC liquid. Following strain separation and the reviving of rotating partners, the next round of selection was initiated following the methods for washing, dilution, and co-culturing of partner strains described above. To minimize positional/edge effects, treatment positions were serially rotated between well positions during each transfer.

### Growth Assays

After 54 weeks of evolution, a single CFU was selected as a representative isolate for each population and used in growth assays. To complete growth assays, co-cultures were prepared as described previously. Controls with the same replicate lineage number (*i.e.* Control A, replicate 1 + Control B, replicate 1) were also co-cultured and are jointly referred to as “Control”. For the LPF treatments, each constantly evolved partner was paired with its most recent rotating partner strain from passage 54 (Rotating Partner A3 and Rotating Partner B3), but growth in these treatments did not vary in response to rotating partner strain (Figure S2). Five microliters of each co-culture were seeded onto the center of SC-Leu-Trp wells (1mL) in a 48-well plate. Colony size was tracked by capturing images of individual wells using the 0.5x objective of a Zeiss Axio Zoom microscope every 24 hours for four days.

We quantified colony size by measuring the two-dimensional area of each colony at each recorded time point. Recent work has demonstrated that this is a reliable method for measuring the growth of *S. cerevisiae* and correlates strongly with measurements of OD600 (Miller et al. 2022). Moreover, measuring growth on agar most accurately reflects the fitness of mutualistic yeast in their selective environment. To minimize positional/edge effects, treatment positions were serially rotated between well positions across replicate assays. We completed five replicate assays for the data shown in Figure 2B-C and four replicate assays for the data shown in Figure 2D-E. For data shown in the supplemental, we completed four replicate assays to quantify ancestral growth rates (Figure S1) and five replicate assays to quantify the effect of each rotating partner on fitness in the LPF-A and LPF-B treatments (Figure S2). Images were assigned random identifiers for blinded analysis and analyzed in ImageJ. We adjusted the threshold of each image using custom settings (based on lighting of image sets), inverted the image, and analyzed particles to identify colonies as regions of interest (ROI). We manually verified correct ROI detection for each image. We used ImageJ’s built-in “Measure” tool to quantify colony area, and treatment identifiers were restored after completed analysis.

**Figure 2.**
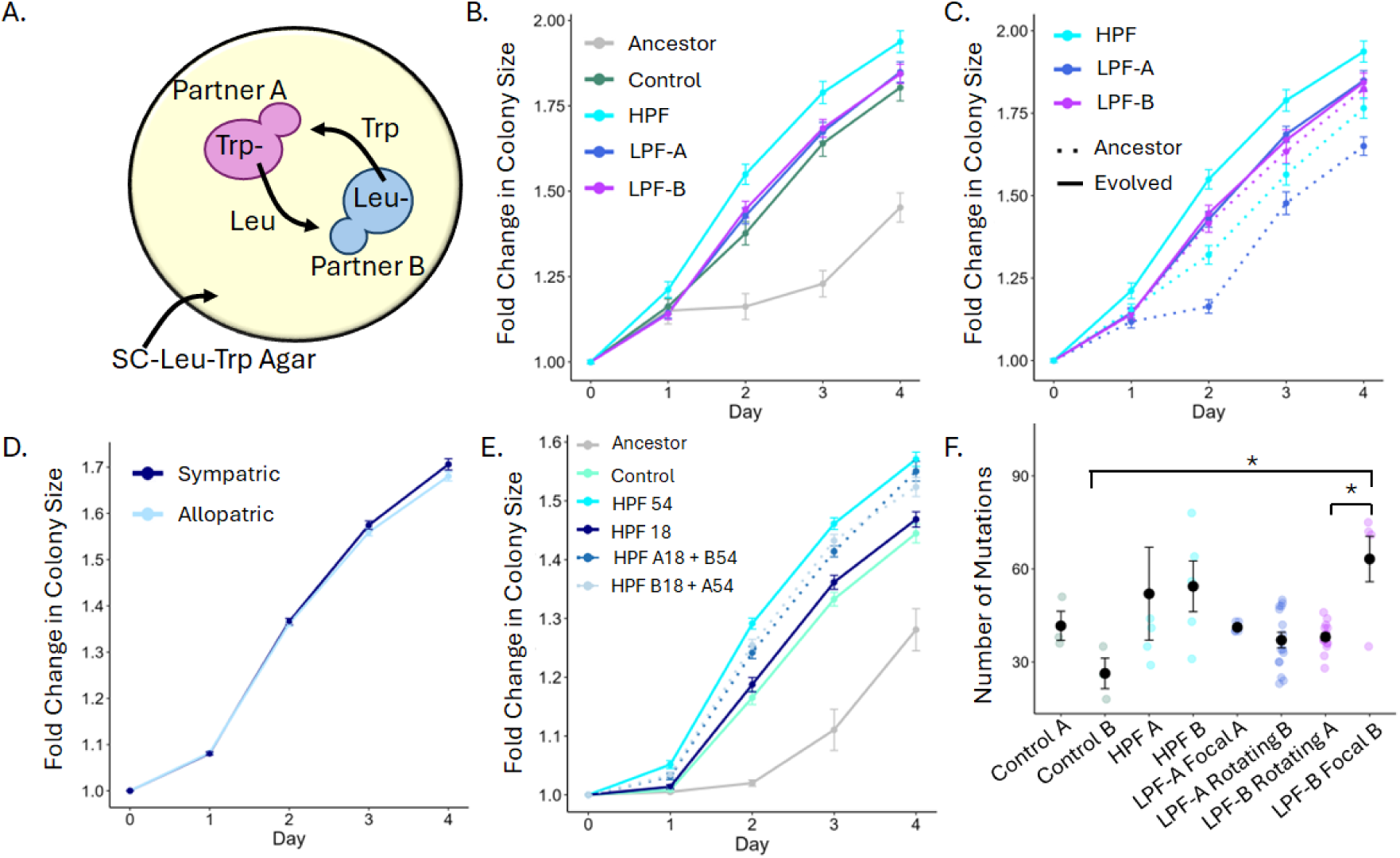
Mutualistic benefits and coevolutionary dynamics vary across treatments. A) Cartoon of metabolic exchange between mutualistic partners. Partner A is auxotrophic for tryptophan (Trp) but overproduces and exports leucine (Leu). Partner B is auxotrophic for Leu but overproduces and exports Trp. Cells grown on Synthetic Complete (SC) agar lacking Leu and Trp (SC-Leu-Trp) are obligately dependent on one another for growth and replication. B) Fold change in colony size across experimental treatments. Points and error bars represent mean fold change and error across replicate lineages. The HPF treatment showed increased growth relative to all other treatments. LPF treatments did not differ from each other, but LPF-B exhibited significantly increased growth compared to control (*t* = 2.84, df = 298, *p* = 0.029), while LPF-A showed only marginally increased growth (*t* = 2.42, df = 298, *p* = 0.096). C) Growth across treatments in time shift assays. Solid lines represent co-cultures between two evolved lineages (passage 54), and dashed lines represent co-cultures between evolved lineages (passage 54) and ancestral partners. In HPF and LPF-A treatments, co-cultures of two evolved partners showed increased growth compared to those with an ancestor. D) Local adaptation assay results showing growth of sympatric (dark blue) versus allopatric (light blue) combinations of HPF lineages from reciprocal inoculations. Selection in HPF treatment did not result in lineage-specific specialization. E) Growth comparison of HPF lineages at passage 54, passage 18, and time-shifted passage 54 + passage 18 co-cultures. Passage 54 co-cultures showed increased growth compared to control, passage 18 co-cultures, and both time-shifted combinations. Both passage 54 + passage 18 co-cultures exhibited increased growth relative to control and passage 18 co-cultures, while passage 18 co-cultures did not differ from control. F) Number of mutations evolved in each treatment. Black points and error bars represent the mean number of mutations and error across replicate lineages. Colored points show the number of mutations for each independent replicate lineage according to treatment. Focal Partner B lineages from the LPF-B treatment exhibited a significantly higher number of mutations relative to their co-passaged rotating A partners (*t* = 3.597, df = 48, *p* = 0.021) and relative to Control B lineages (*t* = 3.73, df = 48, *p* = 0.014). Note for B&C) Growth in each treatment was measured across Days 1-4, and for visualization, we normalized colony area by the mean area of reference colonies at Day 0. Statistics were performed with Day 0 excluded. We later performed separate assays and confirmed there was no effect of treatment on Day 0 area (*F_4,90_* = 0.052, *p* = 0.995).

### Reference colonies to quantify initial colony size

To plot and analyze colony growth, we converted raw two-dimensional area values to fold change in colony size by dividing the area from Days 1-4 by the size of the inoculums on Day 0. We failed to capture Day 0 values for the data shown in Figure 2B&C and Figure S2. To estimate initial colonize size at Day 0 for these assays, we independently seeded and quantified the area of 70 reference cultures. We calculated the mean size of reference cultures (15011.678 mm^2^) and normalized all observed Day 1-4 area values by this mean reference value. To plot growth over time, we replicated the Day 0 values across all treatments so that all treatments have the same starting value, allowing us to visualize the differences between treatments on Days 1-4 without plotting spurious differences between treatments on Day 0. To perform statistical analysis, Day 0 was removed from the dataset. Because all data were normalized by the same value this approach did not alter statistical associations between treatments, and normalization simply improved visualization.

However, to confirm that this approach did not alter statistical associations, we also used raw area (without conversion to fold change) values to assess the effect of treatment on colony size, excluding Day 0, and the results were consistent with our fold change approach (See Figure S3 and Supplemental Data 1). We then also confirmed that there was no effect of treatment on initial colony size that may have driven differences on Days 1-4. To do this, we performed five independent replicate assays where we co-cultured the co-passaged partners from each replicate lineage of each treatment and plated them on SC-Leu-Trp as described above. We measured the area of the colony immediately after plating and used a one-way ANOVA to test for an effect of treatment on initial colony area. We observed no differences between treatments (Figure S4; *F_4,90_* = 0.052, *p* = 0.995). For all other growth assays, we directly examined droplet size at Day 0. For these assays, Day 0 colony size also did not differ between treatments. Importantly, these results demonstrate that initial colony size did not differ between treatments across all assays and confirms that our approach for normalizing the data in Figure 2B&C did not alter our interpretation of the relationships between treatments.

### Genomic DNA preparation and whole genome sequencing

To determine the genetic basis of observed fitness differences between treatments, we performed whole-genome sequencing of the evolved isolates from passage 54 and the ancestral genomes. Yeast strains were revived from glycerol stocks by streaking onto SC agar. Single colonies were grown overnight in YPD [2% dextrose (w/v), 2% peptone (w/v), 1% yeast extract (w/v)], and DNA was extracted from 1mL aliquots using an IBI Scientific gYeast Genomic DNA kit.

Illumina sequencing libraries were prepared by SeqCenter using the tagmentation-based and PCR-based Illumina DNA prep kit and custom IDT 10bp unique dual indices (UDI) with a target insert size of 280bp. Illumina sequencing was performed on an Illumina NovaSeq X Plus sequencer. Paired-end 150bp samples were prepared for each sample.

Hybrid Illumina + Nanopore genomes were prepared for the ancestral genomes by SeqCenter. According to Seqcenter’s protocol, nanopore libraries were prepared using the PCR-free Oxford Nanopore Technologies Ligation Sequencing Kit (SQK-NBD114.24) with the NEBNext Companion Module (E7180L) according to the manufacturer’s specifications. Nanopore sequencing was performed on an Oxford Nanopore a MinION Mk1B sequencer using the 400bps sequencing mode with a minimum read length of 200bp. Adapters were removed from Nanopore reads using Porechop (an open source software for the QC and adapter trimming of ONT technologies n.d.). *De novo* genome assemblies were generated from the Oxford Nanopore Technologies (ONT) read data with Flye2 (Kolmogorov et al. 2019) under the nano-hq (ONT high-quality reads) model. Subsequent polishing used the Illumina read data with Pilon3 (Walker et al. 2014) under default parameters. To reduce assembly artifacts caused by low quality nanopore reads, long read contigs with an average short read coverage of 15x or less were removed from the assembly. Median contig length for Ancestor A was 3253 and for Ancestor B was 3493 and strains had a median coverage across nanopore reads of ∼50 X. Assembled genome sequences were annotated using funannotate (J 2017).

### Sequence analysis

We processed Illumina reads using Trimmomatic v0.39 (Bolger et al. 2014) to remove low-quality bases and adapter sequences. Regions where quality score fell below 20 and reads shorter than 30bp were trimmed to reduce artifacts caused by low quality reads. Relationships between clonal isolates were analyzed using breseq (Deatherage and Barrick 2014), which has been commonly used for experimentally evolved strains of *S. cerevisiae* (Keane et al. 2014; Morard et al. 2019; Pereira et al. 2019; Lairón-Peris et al. 2020, 2021; Barber et al. 2021), and is specifically well-suited for our system since our mutualistic yeast strains are haploid and obligately asexual (Barrick et al. 2014). We prepared a hybrid assembly for the ancestor of each partner genotype, and Illumina reads were aligned against these ancestral genomes of Partner A and Partner B, which served as the reference genomes for breseq analysis. Illumina reads for each ancestral genome were also aligned against the ancestral hybrid assemblies to control for sequencing artifacts. Mutations identified in ancestral short read genomes were filtered from the GenomeDiff “.gd” files (breseq version of a VCF file) of the evolved isolates using gdtools subtract. The resulting .gd files were used to compare the presence/absence, number, and types of mutations across all genomes.

We detected coding regions where replicate lineages exhibited evidence of convergent evolution by filtering the genomes to display only missense and nonsense mutations where more than one lineage evolved a mutation. We confirmed signatures of convergent evolution at specific genes across lineages did not result from cross-contamination by evaluating whether mutations resulted from the same base substitutions. We observed unique mutations arising in genes that convergently evolved mutations across experimental lineages, indicating a lack of cross-contamination (see Supplemental Table 1&2). If mutated genes were also identified in the control treatment, we tested for a significant co-occurrence with other mutated genes in the genome, which may indicate an important epistatic interaction. If no significant co-occurrences were observed, we filtered mutated genes that were also present in the control to identify mutations resulting from treatment-specific selection. We tested for significant co-occurrences between genes using Fisher’s exact test. We considered patterns of parallel genomic evolution across replicate populations as evidence of adaptive change resulting from selection (Nakatsu et al. 1998; Cooper et al. 2001, 2003).

We further examined the distribution of mutations for signatures of selection by simulating a null distribution of 100,000 different types of mutations (missense, synonymous, nonsense, intergenic, and small indel) in the ancestral genomes using the Python package Mutation-Simulator (Kühl et al. 2021). Specifically, we simulated a total of 40,000 small indels and 60,000 SNPs, maintaining the same SNP to small indel ratio as in the observed data. We built custom annotation databases in SnpEff (Cingolani et al. 2012) using our annotated ancestral assemblies and use these databases to annotate the mutations in the simulated genomes. We then used the python package EMUs (Dubose 2023) to run 1000 bootstrap simulations where we sampled the observed number of each mutation type for each treatment (using the sum of each mutation type across all replicate lineages). The observed frequency of each mutation type was compared to the null distribution, which is the expected distribution of mutations under genetic drift. For all treatments, we observed a lower frequency of frameshift mutations (small indels in coding regions) than expected at random small indels in non-coding regions did not differ from expectations at random, so in the main text we focused our analysis on point mutations.

### Statistics

We used the R package ‘lmerTest’ (Kuznetsova et al. 2017)’ to analyze growth across treatments using mixed effects linear models. In all models, we treated replicate lineage and assay replicate as random effects. This R package uses the Satterthwaite approximation to estimate degrees of freedom (due to the hierarchical structure of the model), which produces non-integer values (Kuznetsova et al. 2017). When significant main effects were observed, we used the R package ‘emmeans’ (Lenth 2022) to predict pairwise relationships between treatments using estimated marginal means based on the models. We tested for differences in the number of mutations evovled across experimental treatments using a one-way ANOVA, and we tested whether the frequency of each mutation type (missense, synonymous, nonsense, intergenic, small indel) varied in response to treatment using a generalized linear model. For all posthoc analyses, using estimated marginal means to predict pairwise relationships we applied a Bonferroni correction for repeated tests. Figures were prepared in R using ‘ggplot2’ (Whickam 2016) and data was prepared using ‘dpylr’ (Wickham H, François R, Henry L 2022). All statistics were performed in R version 4.2.2.

## Results

We used experimental evolution of genetically engineered *Saccharomyces cerevisiae* strains to test how the temporal fidelity of mutualistic partners affects mutualism evolution. The system consists of Partner A strains (Leu^+^Trp^-^), which are auxotrophic for tryptophan and both overproduce and export leucine (Müller et al. 2014), and Partner B strains (Leu^-^Trp^+^), which are auxotrophic for leucine and overproduce and export tryptophan (Müller et al. 2014). Both amino acids are required for growth. When grown together on media lacking both amino acids, these strains formed an obligate mutualism through metabolic cross-feeding (Figure 2A). Over 54 rounds of selection, we examined the impact of high vs. low partner fidelity on the evolution of mutualism (following the experimental approach summarized in Figure 1).

### MULTI-PARTNER SELECTION CONSTRAINS THE EVOLUTION OF MUTUALISTIC BENEFITS

Mutualists that evolve with high partner fidelity have more opportunities to exert reciprocal selection on one another, which is expected to facilitate the evolution of increased mutualistic benefits as partners become increasingly specialized to one another (Thompson 1994, 2005; Douglas 1998; Batstone et al. 2020). Therefore, we hypothesized that mutualisms with high partner fidelity would evolve increased benefits relative to those evolved with low partner fidelity. We tested this hypothesis by assessing mutualism fitness at the end of experimental evolution (passage 54). We tracked growth by plating 50:50 inocula of partner strains plated onto solid selective media, then monitoring growth over four days. Fold change in colony size was calculated over time by dividing the colony area from each day by the mean area of reference colonies measured at plating (see methods), and total colony growth was considered the fold change in growth after four days. For the LPF treatments, each focal mutualist was paired with its most recent rotating partner strain from passage 54 (*i.e.,* Rotating Partner A3 and Rotating Partner B3), but growth in these treatments did not vary in response to rotating partner strain (Figure S1).

Using a Linear mixed-effects model (LMM), we observed significant effects of day (LMM: *F_3,355.97_* = 1500.28, *p* < 0.0001), treatment (LMM: *F_3,308.06_* = 51.65, *p* < 0.0001), and a marginal treatment × day interaction (LMM: *F_9,335.97_* = 1.89, *p* = 0.053) on colony growth. The HPF treatment exhibited significantly increased growth compared to the control with 7.2% higher overall growth (Figure 2B; total mean fold change: 1.94 [HPF] vs. 1.81 [Control]; *t* = 10.453, df = 298, *p_adj_* < 0.0001, Cohen’s *d* = 0.85, 95% CI [0.17, 1.55]). Relative to both Low Partner Fidelity treatments (LPF-A and LPF-B), which differ according to whether Partner A or Partner B is the focal strain with associated rotating partners, the HPF treatment also exhibited a significant increase in growth (Figure 2B; HPF vs. LPF-A: *t* = 9.6, df = 336, *p_adj_* < 0.0001; HPF vs. LPF-B: *t* = 9.107, df = 336, *p_adj_* < 0.0001) with a total of 4.9% higher overall growth compared to both LPF-A and LPF-B (Figure 2B; HPF vs. LPF-A: total mean fold change = 1.94 vs. 1.85, Cohen’s *d* = 0.57, 95% CI [-0.01, 1.15]; HPF vs. LPF-B: total mean fold change = 1.94 vs. 1.85, Cohen’s *d* = 0.63, 95% CI [0.04, 1.21]).

LPF-A and LPF-B had similar growth rates (Figure 2B; *t* = 0.497, df = 336, *p_adj_* = 1), and the same mean fold change in growth after four days (Figure 2B; LPF-A = 1.85 ± 0.151, LPF-B = 1.85 ± 0.140). Relative to Control, LPF-B exhibited significantly elevated growth on average (Figure 2B; *t* = 2.84, df = 298, *p_adj_* = 0.029), while LPF-A showed only a marginal increase in growth on average (Figure 2B; *t* = 2.42, df = 298, *p_adj_* = 0.096). However, both treatments had similar small effect sizes on total colony growth, with each resulting in only a 2.2% increase in total growth compared to Control (LPF-A: Cohen’s *d* = 0.30, 95% CI [-0.36, 0.96]; LPF-B: Cohen’s *d* = 0.28, 95% CI [-0.39, 0.94]). These results demonstrate that increased mutualistic benefits evolved most rapidly in mutualisms with high partner fidelity. Despite evolving under selection for mutualism, the LPF treatments exhibited proportionally lower increases in mutualistic benefits, with only modest benefits over the control. This demonstrates that generalism does not necessarily inhibit the evolution of mutualistic benefits, but it does slow rates of adaptation, likely due to the limited coevolutionary potential between partners.

### COEVOLUTION IS NOT REQUIRED FOR THE RAPID EVOLUTION OF MUTUALISTIC REWARDS

We hypothesized that rapid evolution of increased mutualistic rewards in the HPF treatment resulted from the tighter coevolutionary dynamics made possible by consistent partner pairings, compared to the more limited coevolutionary potential in the LPF treatments. We tested this hypothesis using time shifts (Brockhurst and Koskella 2013) to compare the growth of co-cultures between evolved isolates at passage 54 (*e.g.*, HPF-A.1-54 + HPF-B.1-54) relative to co-cultures between evolved isolates and an ancestral partner (*e.g.* HPF-A.1-54 + ANCB). For time shifts in the LPF treatment, focal mutualists were paired with their most recent partner (*i.e.*, A3 or B3) from passage 54. If mutualistic benefits resulted through coevolution, we expected co-cultures between two evolved isolates to exhibit increased colony growth relative to co-cultures involving an ancestor (Brockhurst and Koskella 2013; Vidal and Segraves 2021).

We observed significant effects of day (LMM: *F_3, 588.01_* = 1504.89, *p* < 0.0001), treatment (LMM: *F_2, 588.03_*, *p* < 0.0001), partner (LMM: *F_1, 589.23_* = 232.72, *p* < 0.0001), treatment x day (LMM: *F_6,588.03_* = 3.97, *p* < 0.0007), partner x day (LMM: *F_3, 588.01_* = 18.71, *p* < 0.0001), and treatment x partner x day (LMM: *F_6,588.01_* = 3.07, *p* < 0.0001) on colony growth. Overall, treatment and partner substantially impacted colony growth, with partner effects dependent on treatment (Figure 2C). For example, HPF lineages had increased fitness when co-cultured with their evolved partners compared to an ancestral partner (Figure 2C; *t* = 13.52, df = 597, *p_adj_* < 0.0001), resulting in a 10% relative increase in overall growth (Cohen’s *d* = 0.95, 95% CI [0.41, 1.49]), and consistent with coevolution.

We next examined evidence for coevolution within each LPF treatment. In the LPF-A treatment, mutualistic growth was increased when Focal Partner A was paired with its evolved Rotating Partner B3 relative to Ancestor B, with 12% higher overall growth (Figure 2C; *t* 10.32, df = 596, *p_adj_* < 0.0001, Cohen’s *d* = 1.42, 95% CI [0.74, 2.10]). This too is consistent with coevolution. In contrast, in the LPF-B treatment, mutualism fitness did not differ when Focal Partner B was paired with Ancestor A versus its evolved Rotating Partner A3 (Figure 2C; *t* = 1.29, df = 597, *p_adj_* = 1). This suggests that the increase in mutualistic benefits observed in the LPF-B treatment was driven by the one-sided evolution of the Focal Partner B, rather than through coevolution. Interestingly, Ancestor Partner A exhibited a significantly higher baseline replication rate than Ancestor Partner B (Figure S2), suggesting that whether coevolution occurs may depend on the growth rate of the focal partner. Specifically, coevolution may be impeded by slow adaptation in the ‘weakest link’ partner, whose growth rate is limiting to the mutualism. Moreover, these findings suggest that neither partner fidelity nor coevolution is strictly necessary for the evolution of increased mutualistic benefits, although the strongest benefits emerged in the HPF treatment where partner fidelity was high and coevolution likely occurred.

### HIGH PARTNER FIDELITY RESULTS IN INCREASED MUTUALISTIC BENEFIT WITHOUT PARTNER SPECIFICITY

Evidence from the preceding time shifts suggests that partner fidelity and coevolution jointly contributed to the rapid evolution of increased mutualistic benefits in the HPF treatment. Reciprocal selection between mutualistic partners that increases their specialization to one another can facilitate the evolution of mutualistic benefits (Thompson 1994, 2005; Douglas 1998; West et al. 2015). Therefore, we hypothesized that the rapid evolution of mutualistic benefits in the HPF treatment resulted from the reciprocal specialization of co-passaged lineages. To test this, we used reciprocal cross inoculations to examine whether co-passaged strains isolated from the same replicate population were locally adapted to one another (5 Partner A lineages x 5 Partner B lineages = 25 cross inoculations). Local adaptation is considered a signature of specialization between coevolved lineages (Kawecki and Ebert 2004; Blanquart et al. 2013; Brockhurst and Koskella 2013), and we expected to observe increased fitness between sympatric partners relative to allopatric partners if this specialization evolved (Petit and Thompson 1998; Kawecki and Ebert 2004; Blanquart et al. 2013).

Using a linear mixed effects model, we tested for a significant Partner A x Partner B interaction on colony growth, which is expected if strains are locally adapted to one another (Blanquart et al. 2013). We observed a significant Partner A x Partner B interaction (LMM: *F*_16_*_,467_* = 2.14, *p* = 0.006) and used a linear contrast to test whether this interaction resulted from differences between sympatric and allopatric partner combinations. The contrast revealed no difference between sympatric and allopatric combinations (*t* = 0.382, df = 471, *p* = 0.70), indicating a lack of local adaptation (Figure 2D). This suggests that genetic specificity did not evolve between co-passaged lineages (Blanquart et al. 2013).

We observed no evidence for specialization between co-passaged HPF lineages, yet evolved mutualists exhibited increased fitness when paired with one another relative to when paired with the ancestor of their co-passaged partner. This pattern suggests co-adaptation occurred but was not partner-specific, leading us to propose two alternative explanations. First, in the HPF treatment both partners experienced prolonged exposure to selection relative to the rotating partners in the LPF treatments, and increased benefits may have resulted from prolonged parallel adaptation to a shared environment rather than through coevolution (Janzen 1980; Thompson 1994; Antonovics et al. 2013). Alternatively, the rapid evolution of mutualistic benefits may have resulted through coevolution without detectable partner specificity if replicate populations followed similar evolutionary trajectories, with resource sharing generally improving over time, similar to coevolutionary arms race.

Performing additional time shifts for the HPF treatment allowed us to further evaluate the temporal dynamics between partners and gain greater insights into whether coevolution played a role in these mutualisms despite the lack of genetic specificity between co-passaged lineages. We performed time shifts using the following co-cultures: Passage 18A + Passage 18B, Passage 54A + Passage 54B, Passage 54A + Passage 18B, Passage 54B + Passage 18A, and Control A + Control B. If coevolution facilitated the evolution of mutualistic benefits observed in the HPF treatment, we expected mutualists from future time points (passage 54) would be well adapted to their partners from past time points (passage 18), and we expected time-shifted co-cultures would produce increased growth relative to both control and passage 18 co-cultures, although constrained relative to passage 54 co-cultures. If adaptation to the environment drove increased fitness without reciprocal adaptation, we expected passage 54 co-cultures and time-shifted co-cultures to exhibit similar growth supported by the adaptation of passage 54 isolates alone.

We observed a significant effect of partner (LMM: *F_4,427.92_* = 76.20, *p* < 0.0001), day (LMM: *F_4,426.91_* = 3387.69, *p <* 0.0001), and partner × day (LMM: *F_16,426.91_* = 8.19, *p* < 0.0001) on colony growth (Figure 2E). Passage 54 co-cultures exhibited significantly increased growth on average compared to Control (*t* = 14.40, df = 431, *p_adj_* < 0.0001), Passage 54A + Passage 18B (*t* = 5.129, df = 427, *p_adj_* < 0.0001), Passage 54B + Passage 18A (*t* = 5.40, df = 427, *p_adj_* < 0.0001), and Passage 18 co-cultures (*t* = 13.42, df = 427, *p_adj_* < 0.0001). Overall, Passage 54 treatments exhibited increased total colony growth of 3.3% relative to Passage 54A + Passage 18B (Cohen’s *d* = 0.74, 95% CI [0.07, 1.42]), 1.3% relative to Passage 54B + Passage 18A (Cohen’s *d* = 0.31, 95% CI [-0.34, 0.97]), and 6.8% relative to passage 18 co-cultures (Cohen’s *d* = 1.84, 95% CI [1.06, 2.61]). Time-shifted co-cultures where Partner B was ahead of Partner A in evolutionary time (e.g., Passage 54B + Passage 18A) were more similar to Passage 54 co-cultures than time-shifted co-cultures where Partner A was ahead of Partner B (e.g., Passage 54A + Passage 18B). This indicates that fitness in the HPF treatment depended more heavily on the evolution of Partner B, which exhibits a slower baseline replication rate than Partner A (Figure S2).

Both time-shifted co-cultures (Passage 54A + Passage 18B and Passage 54B + Passage 18A) showed intermediate growth rates that were significantly higher than both the control (P54A+P18B: *t* = 10.11, df = 431, *p* < 0.0001; P54B+P18A: *t* = 8.84, df = 431, *p_adj_* < 0.0001) and Passage 18 co-cultures (P54A+P18B: *t* = 8.31, df = 427, *p_adj_* < 0.0001; P54B+18A: *t* = 8.04, df = 427, *p_adj_* < 0.0001) but lower than Passage 54 co-cultures. Time-shifted co-cultures had large effects on growth relative to Passage 18 co-cultures with an increase in total colony growth of 3.4% for Passage 54A + Passage 18B (Cohen’s *d* = 0.84, 95% CI [0.18, 1.51]) and 5.4% for Passage 54B + Passage 18A (Cohen’s *d* = 1.22, 95% CI [0.53, 1.92]) (Figure 2E). These dynamics are consistent with coevolution, where reciprocal selection between partners likely favored an increase in the exchange of mutualistic resources through time. Importantly, the growth of the time-shifted co-cultures (Passage 54 A + Passage 18B and Passage 54B + Passage 18A) was consistently elevated relative to Control, demonstrating that HPF lineages exhibited increased fitness even when co-cultured partners experienced asymmetric exposure to selection. Therefore, the improved fitness in the HPF treatment relative to the LPF-A and LPF-B treatments did not arise solely from mutualists in the HPF treatment experiencing prolonged parallel adaptation to a shared environment.

### SIGNATURES OF SELECTION IN THE GENOMES OF BOTH PARTNERS

To investigate the genetic basis of the divergent evolutionary outcomes across treatments, we performed whole genome sequencing of a single representative isolate from each replicate population and detected evolved mutations using breseq (Deatherage and Barrick 2014), which has been routinely used for genomic analysis of *S. cerevisiae* (Keane et al. 2014; Morard et al. 2019; Pereira et al. 2019; Lairón-Peris et al. 2020, 2021; Barber et al. 2021). We observed a significant effect of treatment (*F_7,287_* = 7.37, *p* < 0.0001), mutation type (*F_5,287_* = 286.77, *p* < 0.0001), and treatment x mutation type (*F_35,287_* = 5.11, *p* < 0.0001) on the number of evolved mutations. The number of mutations evolved in Control A and Control B did not significantly differ from one another (*t* = 1.388, df = 48, *p_adj_* = 1), suggesting that in the absence of selection for mutualism, Partner A and Parter B have similar rates of mutation accumulation. However, in treatments where obligate mutualistic cross-feeding was required, we observed an increase in mutation number for Focal B lineages in the LPF-B treatment relative to their co-passaged Rotating A Partners (*t* = 3.597, df = 48, *p* = 0.02) and Control B (*t* = 3.73, df = 48, *p* = 0.014) (Figure 2F). The number of evolved mutations was similar between co-passaged partners in all other treatments. This suggests that Focal B lineages in the LPF-B treatment were under particularly strong selection, further indicating that evolution of mutualistic benefits in this treatment was driven primarily by Focal Partner B adaptation.

We then compared the differences between treatments for each mutation type. We observed a significantly greater number of small indels for HPF-A lineages relative to HPF-B lineages (*t* = 3.536, df = 287, *p* = 0.01). Within the LPF-A treatment, Focal Partner A lineages exhibited a greater number of intergenic mutations than their co-passage Rotating Partner B (*t* = 3.937, df = 287, *p* = 0.0029). Finally, HPF Partner B and LPF-B Focal Partner B lineages evolved more missense mutations than their co-passaged Partner A lineages (HPF-B vs. HPF-A: *t* = 5.30, df = 287, *p* < 0.0001; LPF-B Focal Partner B vs. Rotating

Partner A: *t* = 9.744, df = 287, *p* < 0.0001) (Figure 3A&B). Overall, treatments exhibited differences in the types of mutations they evolved, indicating that selection acted differently on lineages of Partner A (more intergenic mutations and small indels) and Partner B (more mutations in coding regions).

**Figure 3.**
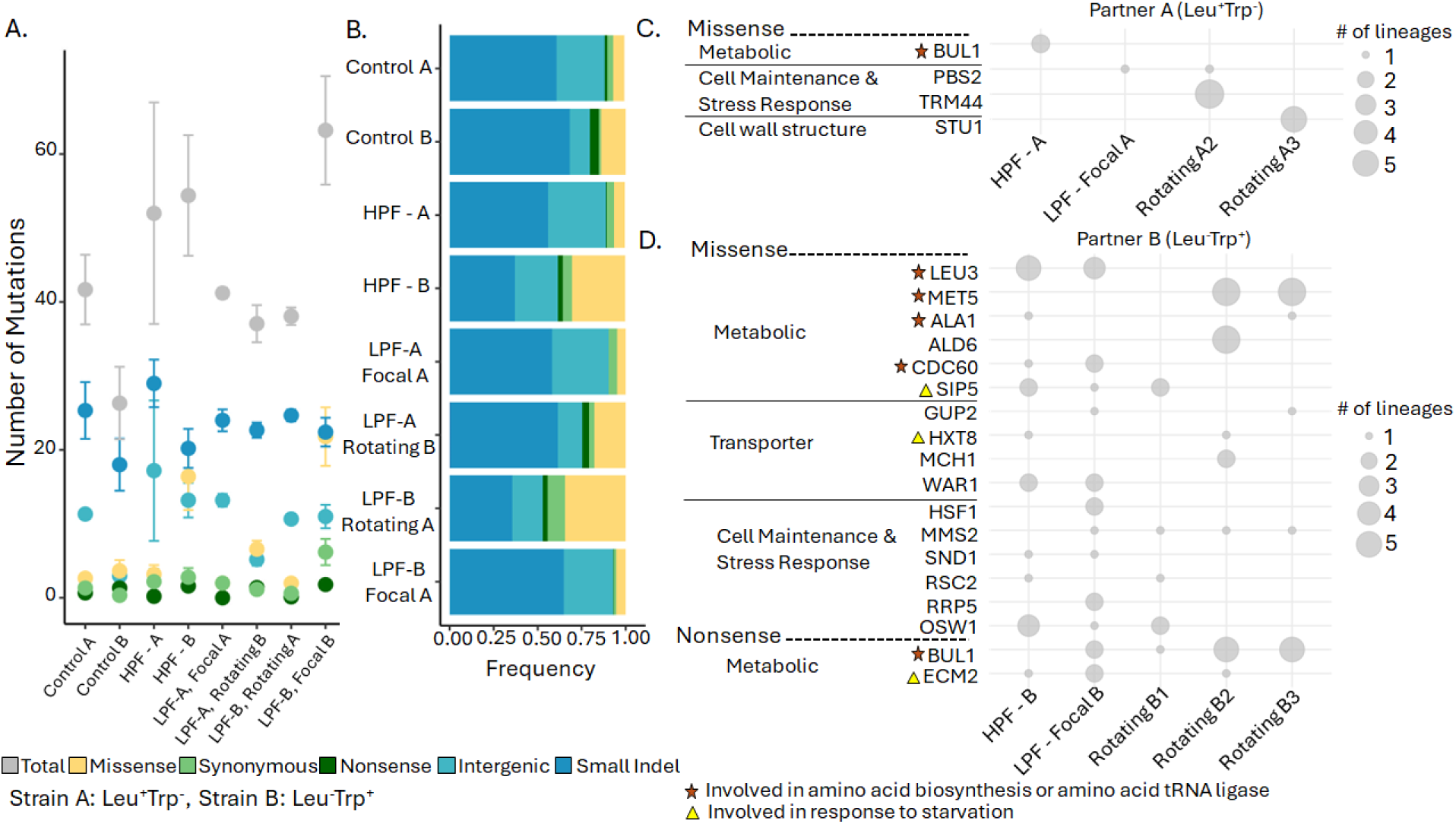
Partner B lineages show signatures of convergent evolution across coding regions. A) Number of each mutation type observed in each treatment. Points represent the mean number of mutations across replicate lineages, and error bars show the standard error across replicate lineages. Partner B lineages evolved a significantly greater number of missense mutations than Partner A lineages. B) Relative frequencies of mutation types within each treatment. Frequencies of missense mutations were elevated in Partner B lineages. C) Convergent evolution in Partner A coding regions. Partner A lineages convergently evolved fewer mutations in coding regions than Partner B lineages. Lineages of Rotating A1 exhibited no evidence of convergent evolution with any other Partner A lineages. D) Convergent evolution of coding regions across Partner B lineages. Partner B convergently evolved mutations in 18 genes, including 5 genes involved in amino acid biosynthesis or amino acid tRNA ligation, and 3 genes involved in responses to starvation. C&D) Genes that also evolved mutations in the controls were filtered, unless they shared a significant co-occurrence (determined using Fisher’s exact test) with other mutated genes. Dot size represents the number of replicate lineages with mutations in the specified gene. Orange star indicates a gene is involved in amino acid biosynthesis or ligation of amino acids to tRNA. Yellow triangle indicates a gene is involved in stress response to starvation.

Parallel genomic evolution across replicate populations is generally recognized as evidence of adaptive change resulting from selection (Nakatsu et al. 1998; Cooper et al. 2001, 2003). Therefore, to further evaluate evidence for adaptive evolution, we analyzed the distribution of mutations across replicate lineages within each treatment (Figure 3). Partner A lineages generally exhibited little evidence of parallel evolution at coding loci (Figure 3C). We observed only four genes where mutations convergently evolved, and the number of lineages exhibiting convergent evolution was low. We observed parallel evolution of two HPF Partner A lineages that evolved mutations in *BUL1*, involved in the regulation of an amino acid transporter. In contrast, Partner B lineages convergently evolved missense and nonsense mutations in 18 different genes (Figure 3D). Five of these genes were associated with amino acid biosynthesis or amino acid tRNA ligation (*LEU3, MET5, ALA1, CDC60,* and *BUL1*), and three genes were associated with starvation response (*SIP5, HXT8,* and *ECM21*). This genomic signature of parallel evolution indicates that these loci were likely under selection, strongly suggesting that adaptation in each treatment depended heavily on the evolution of Partner B.

Finally, we used a simulation approach to assess the probability that selection rather than neutral processes underpinned the distribution of mutations along the genomes of the evolved lineages within each experimental treatment. To do this, we simulated a null distribution of 100,000 different types of mutations (missense, synonymous, nonsense, intergenic, and small indel) in the Ancestor A and Ancestor B genomes using the Python package Mutation-Simulator (Kühl et al. 2021). We then used the Python Package EMUs (Dubose 2023; Pentz et al. 2023) to run 1000 bootstrap simulations where we randomly sampled the number of observed mutations for each treatment to generate a null distribution of the mutations expected under drift. We compared the number of observed mutations within each treatment to this null distribution (Figure S5). When the number of mutations of a given type (e.g., nonsynonymous) was unlikely given how often these mutations arise by chance (i.e., in the lower 2.5% or upper 97.5% of simulated values in our null model), we considered this as evidence of selection. Our simulations suggest that selection acted on both Partner A and Partner B lineages across several treatments. Missense mutations occurred significantly less in the observed data than expected by chance for Focal Partner A lineages in both the HPF and LPF-A treatments and for Rotating Partner A lineages in the LPF-B treatment (observed values occurred less than 2.5% of the simulated values) (Figure S5). This suggests that coding loci in Partner A lineages were under purifying selection. In contrast, for replicate lineages of Rotating Partner B in the LPF-A treatment, intergenic mutations in the observed data occurred with marginally increased probability than expected by chance (observed value was greater than 97% of simulated values; Figure S5), suggesting these lineages were under positive selection. For lineages of Focal Partner B in the LPF-B treatment, missense mutations were marginally increased relative to expected at random occurring at greater than 97% of simulated values (Figure S5). Combined with signatures of parallel evolution (Figure 3), this suggests that some Focal Partner B lineages in the LPF-B treatment were also under positive selection.

## Discussion

Whether mutualisms involve few or many partners can depend on ecological dynamics that alter the frequency of interaction between lineages across space or time (Thompson 1994; Batstone et al. 2018; Chomicki et al. 2020; Vidal et al. 2020, 2024). Conditions that facilitate repeated interactions between specific beneficial pairs of lineages increase opportunities for specialized interactions through coevolution (Thompson 1994, 2005; Batstone et al. 2020). In contrast, certain ecological conditions, such as asymmetries in the geographic or abiotic ranges of interacting mutualists may limit opportunities for sustained interactions between specific partners, favoring generalism (Wernegreen 2012; Batstone et al. 2018; Chomicki et al. 2020; Vidal et al. 2020; Breusing et al. 2022; Stoy et al. 2023a). For example, *Anasa tristis* squash bugs can acquire and derive the same level of benefit from geographically diverged bacterial symbionts, which possibly facilitates the highly migratory behavior and broad geographic distribution of these insects (Eiben 2004; Stoy et al. 2023a). Moreover, generalist leguminous plant species capable of interacting with diverse rhizobia partners exhibit broader geographic distributions than more specialized legume species (Harrison et al. 2018). How ecologically rich interactions like these shape the evolution of mutualistic benefits and alter opportunities for coevolution that can reinforce cooperation is not well understood.

We directly tested whether partner fidelity alters the evolution of mutualistic benefits and opportunities for coevolution. To do this, we imposed temporal constraints on partner fidelity in the LPF treatments, inhibiting opportunities for specialization. While constrained relative to the HPF treatment, we observed the rapid evolution of increased mutualistic benefit in the LPF-B treatment and marginal increases in the LPF-A treatment in as little as 54 rounds of selection. The evolution of increased benefit in the low partner fidelity (LPF) treatments is particularly pronounced when considering that each rotating partner only experienced 18 total rounds of selection, and these auxotrophic mutualists exhibited very slow initial growth rates (ancestral two-dimensional colony size less than doubled after four days of growth; Figure 2B). Overall, these results demonstrate that mutualistic benefits evolve most rapidly when partners coevolve with high fidelity, but increased mutualistic benefits can rapidly evolve without coevolution even in environmental conditions that preclude specialization.

Surprisingly, we did not observe evidence of localized specialization between co-passaged HPF partners from the same population. We hypothesized several pathways by which these unexpected generalist dynamics in the HPF treatment may have evolved. First, reciprocal adaptation may have occurred within populations without genetic divergence across replicate populations, impeding local adaptation. This may have occurred if there was a narrow evolutionary pathway populations could follow toward increased mutualistic benefits. The simple mechanistic basis of exchanging resources through the environment makes this a likely outcome. While our experimental design required physical interaction between partners for resource exchange (by growing on an agar surface and exchanging resources with neighbors), the mechanism of resource exchange may have limited opportunities for specialization through partner recognition, making divergence in interaction mechanisms across populations unlikely.

Second, specialization may not have evolved if increased mutualistic benefits were driven by the one-sided evolution of a single partner without coevolution. However, when we performed time shifts assays, both the Passage 18A + 54B and Passage 18B + 54A co-cultures exhibited increased growth relative to the Passage 18 co-cultures, and both time-shifted co-cultures exhibited lower fitness than Passage 54 co-cultures (Figure 3E), indicating that increased mutualistic benefits depended on the reciprocal adaptation of both partners. If coevolution occurred within these interactions, it likely resulted through a type of coevolutionary arms race where mutualistic benefit continually ratcheted upward. In this study, we tested for coevolution by measuring the overall fitness of each mutualism and assessed genomic signatures of selection using representative isolates. In the future, we plan to examine the temporal distribution of the relative fitness, costs, and benefits for each partner independently (Vidal and Segraves 2021) and perform population genomics analysis, which may further illuminate underlying coevolutionary dynamics.

Interestingly, we observed distinct outcomes across the two low partner fidelity (LPF) treatments. While both LPF-A and LPF-B evolved similar increases in mutualistic benefits, these benefits evolved through divergent underlying coevolutionary dynamics. We observed dynamics consistent with coevolution in the LPF-A treatment, where the slow-growing Partner B experienced infrequent exposure to selection. However, we did not observe evidence for coevolution in the LPF-B treatment, where the slow-growing Partner B experienced constant exposure to selection (Figure 2B-C). These distinct outcomes may be driven by the differences in growth rate between the co-passaged partners. Within species interactions, partners with the faster generation time are often predicted to be better adapted to their partners (Price 1980; reviewed in Greischar and Koskella 2007). While this may occur in some interactions (Batstone et al. 2020), both theoretical and empirical work have demonstrated this is not a general rule (Lively 1999; Gandon and Michalakis 2002; Greischar and Koskella 2007). The Red King theory of mutualistic evolution even predicts that rates of mutualistic evolution are driven by partners with slower generation times (Bergstrom and Lachmann 2003).

In our LPF-A treatment, slow-growing Partner B strains were rotated temporally but were well-adapted to Focal Partner A (Figure 2C Figure S1). In contrast, in the LPF-B treatment, the fast-growing Partner A strains were rotated temporally and exhibited no evidence of increased adaptation to Focal Partner B relative to Ancestor A (Figure 2C, Figure S1). Consistent with this, in the HPF treatment we observed higher mutualistic fitness in time-shifted co-cultures where Partner B was ahead of Partner A (Passage 54B + Passage 18A) in evolutionary time than for those where Partner A was ahead of Partner B (Passage 54A + Passage 18B), suggesting rates of mutualistic benefits in the HPF treatment depended more heavily on the slow-growing Partner B. Across all treatments, we generally observed dynamics that suggest the (co)evolution of mutualism depends on the slow-growing partner, which can be well-adapted to its fast-growing partner even when partner fidelity is low. This observation is consistent with the Red King Hypothesis, which predicts that rates of evolution within mutualisms are governed by the slow-growing partner (Bergstrom and Lachmann 2003), suggesting this system may be ideal for directly testing this model in future work.

Through experimental coevolution, we demonstrated that mutualistic benefits can evolve rapidly regardless of partner fidelity. By directly comparing high-fidelity pairwise coevolution with diffuse, multi-partner interactions, we isolated the effects of partner fidelity on mutualistic evolution. HPF lineages demonstrated the highest fitness, confirming that partner fidelity confers the most substantial fitness benefits to mutualists. However, mutualistic benefits increased even under low partner fidelity conditions that limited opportunities for tight coevolution. Most remarkably, while mutualistic benefits evolved nearly identically in the LPF-A and LPF-B treatments, the underlying evolutionary dynamics were distinct. LPF-A showed evidence for coevolution while LPF-B did not. This demonstrates that mutualistic evolution can readily occur without reciprocal adaptation, challenging a common assumption that coevolution necessarily plays a central role in mutualism and highlighting the need to directly quantify rather than infer coevolutionary processes. Overall, our results demonstrate that increased mutualistic benefits can rapidly evolve through multiple evolutionary pathways rather than through a single path toward increased specialization, contributing an explanation for the widespread occurrence persistence of generalism.

## Author contributions

Experiments were designed by KSS and WCR. Experimental evolution was conducted by KSS and DL, and experimental assays were completed by KSS. Genomics, data analysis, and statistics were performed by KSS. KSS wrote the initial draft of the manuscript, and all authors contributed to the writing of the final version.

## Acknowledgements

We would like to thank Andrew Murray for generously sharing genetically engineered mutualistic yeast strains. We are also grateful to Mandy Gibson and Kim Hoang for manuscript comments. We also thank Kim Hoang and Ozan Bozdag for helpful conversations about directions and methods for genomics. Finally, we are thankful to Levi Morran and the Ratcliff lab for helpful comments during early data curation.

