## Supplemental Figures and Data for "Partner fidelity and coevolution: Useful but not required for rapidly increased mutualistic benefits"

Supplemental Data and Figures

1)

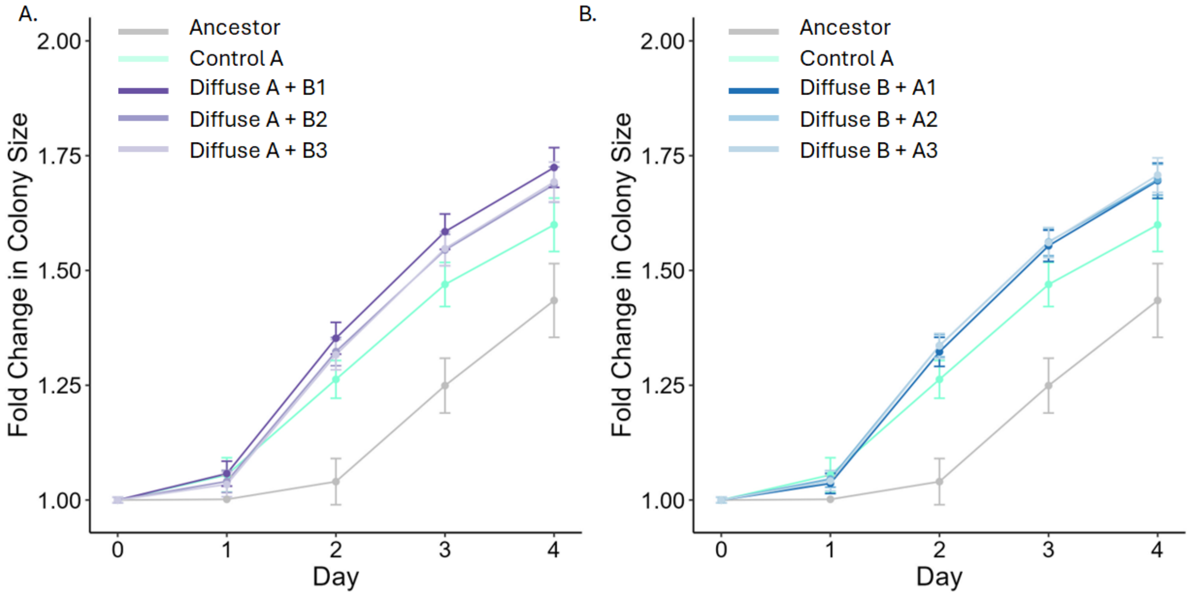


**Figure S1. Identity of rotating partner does not alter fitness in Diffuse treatments.** A) Fitness in the Diffuse A treatment when constantly evolved Diffuse A is paired with each Rotating Partner B. We observed no differences in fitness across partner combinations ($F_{3,160.65}$ = 0.22, $p$ = 0.88). B) Fitness in the Diffuse B treatment when constantly evolved Diffuse B is paired with each Rotating Partner A. We observed no differences in fitness across partner combinations ($F_{3,159.62}$ = 0.19, $p$ = 0.91).

2)

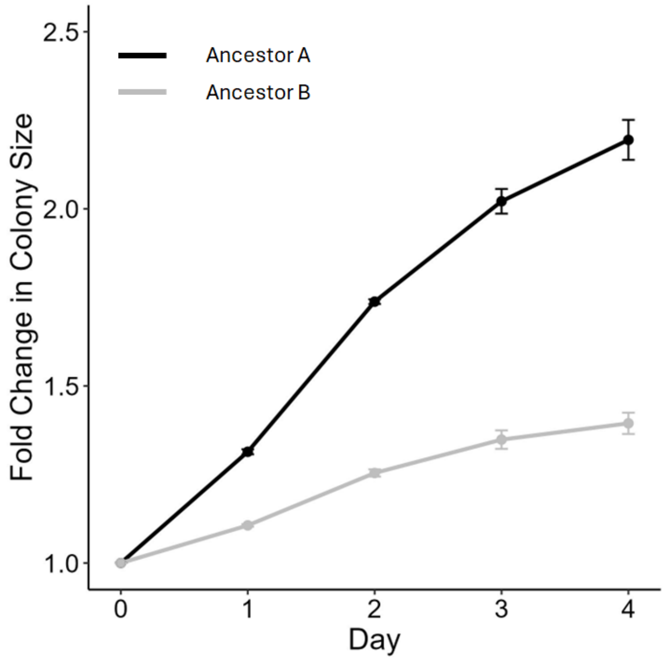


**Figure S2. Generation time of ancestral genotypes.** Ancestor A ($Leu^+Trp^-$) has a significantly faster growth rate than Ancestor B ($Leu^-Trp^+$) **(**LMM: $F = 193.37$, $df = 1$, $p < 0.0001$). Black line shows Ancestor A, and Grey line shows Ancestor B. Points show mean fold change and error across independent assay replicates.

3)

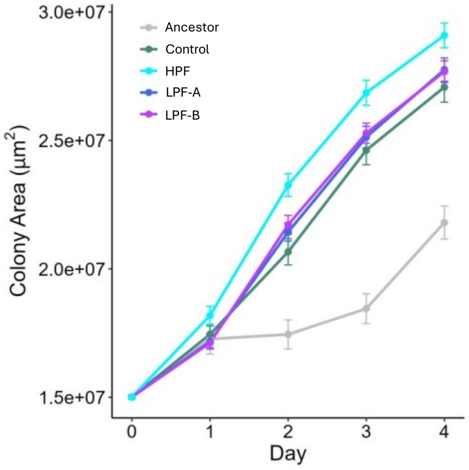


**Figure S3. Change in two-dimensional colony area over four days of growth**. Plot depicts the raw 2D area of growth for colonies in each treatment without conversion to fold change in colony size. Relationships between treatments were not altered by normalizing data by the size of the reference colonies at Day 0 (see supplemental data 2 for statistics.

4)

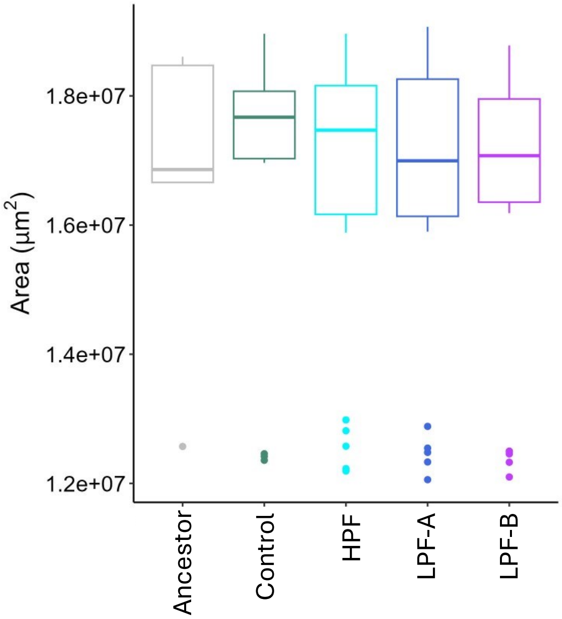


**Figure S4. Day 0 colony size.** We performed five replicate assays to compare the initial colony size of co-cultures from each treatment. We observed no effect of treatment on initial colony size ($F_{4,90}$ = 0.052, $p$ = 0.995). Differences in growth trends across Days 1-4 were not driven by differences in initial colony size.

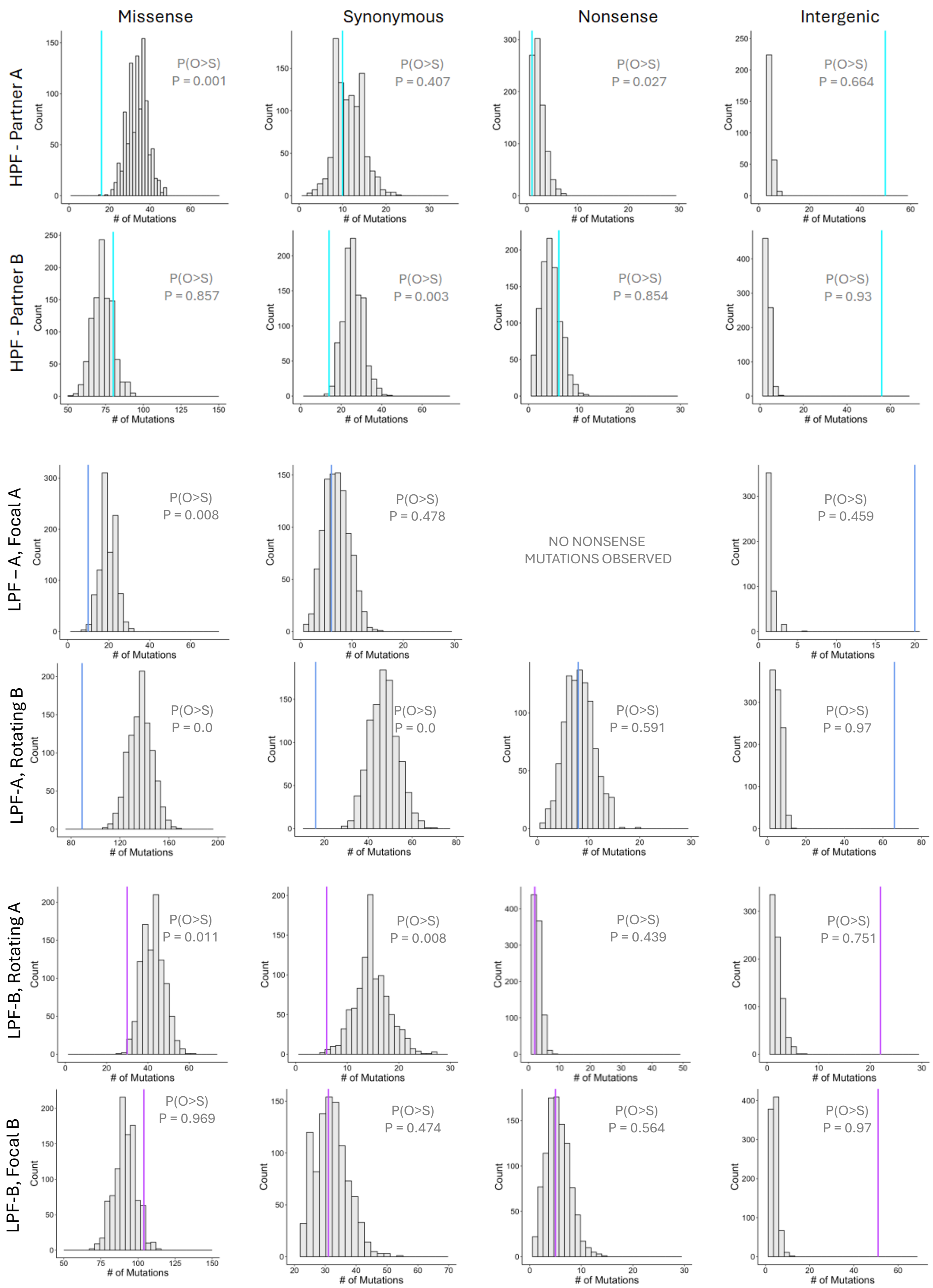


**Figure S5. Simulations reveal signatures of selection on slow-growing Partner B lineages.** P(O>S) is the probability that the number of observed mutations of each type was greater than in those simulated. HPF B and LPF-B Focal B lineages exhibited a higher probability of evolving missense mutations than expected by chance, and HPF B exhibited a higher probability of evolving nonsense mutations than expected by chance, providing evidence for positive selection. Partner and Parter B lineages across all treatments exhibited a higher probability of intergenic mutations, suggesting responses to selection by modulating gene expression.

6) **Supplemental Data 1**. We used a mixed effects linear model to test whether colony area was dependent on the following fixed effects: treatment, day, treatment x day. We included lineage and replicate assays as random effects. Day 0 colony areas were collected by individually plating and quantifying the area of 70 reference colonies. These values were not included in the model. We performed this analysis to confirm that the relationship between treatments was not altered by normalizing the growth data from Days 1-4 by the mean area of reference colonies at Day 0. Normalizing the data by the mean of the reference colonies at Day 0 did not alter relationships between treatments across Days 1-4, and we see the same statistical relationships between treatments in this analysis using raw areas as the model using normalized values which was reported in the main text.

Model results:

| | Sum Sq | Mean Sq | NumDF | DenDF | F-value | p-value |
|---|---|---|---|---|---|---|
| **Treatment** | 1.8281e+14 | 6.0936e+13 | 3 | 2477 | 51.6492 | <2e-16 |
| **Day** | 5.3101e+15 | 1.7700e+15 | 3 | Inf* | 1500.2758 | <2e-16 |
| **Treatment*Day** | 2.0021e+13 | 2.2245e+12 | 9 | Inf* | 1.8855 | 0.0492 |

*Infinite denominator degrees of freedom (Inf) occur when the Satterthwaite approximation estimates a near-infinite number of degrees of freedom. This is common for within-subject/repeated-measures terms with many residual observations and does not affect the validity of the p-values.

Posthoc analysis using emmeans:

| Contrast | Estimate | SE | Df | t ratio | p-value |
|---|---|---|---|---|---|
| Control – LPF-A | -444844 | 184000 | 298 | -2.422 | 0.0962 |
| Control – LPF-B | -521122 | 184000 | 298 | -2.837 | 0.0292 |
| Control-HPF | -1920025 | 184000 | 298 | -10.453 | <.0001 |
| LPF-A-LPF-B | -76277 | 154000 | 336 | -0.497 | 1.0000 |
| LPF-A-HPF | -1475181 | 154000 | 336 | -9.603 | <.0001 |
| LPF-B-HPF | -1398903 | 154000 | 336 | -9.107 | <.0001 |
